# Sharing cells with *Wolbachia*: the transovarian vertical transmission of *Culex pipiens* densovirus

**DOI:** 10.1101/500173

**Authors:** Mine Altinli, Julien Soms, Marc Ravallec, Fabienne Justy, Manon Bonneau, Mylene Weill, Anne-Sophie Gosselin-Grenet, Mathieu Sicard

## Abstract

**Originality-Significance Statement:** This contribution provides a window into the study of a densovirus called CpDV infecting *Culex pipiens* mosquitoes. The originality of the study lies in the fact that CpDV, a DNA virus, was to date almost unstudied, its interaction with its hosts and its transmission mode were unknown. Considering that all *Cx. pipiens* are infected with *Wolbachia,* we studied the influence of this bacterial endosymbiont on CpDV density and transmission. Although *Wolbachia-RNA* virus interactions, especially in non-native *Wolbachia-host* associations have been widely studied, *Wolbachia-DNA* virus interactions were rarely investigated.

In this natural tripartite *Cx. pipiens-Wolbachia*-CpDV system:

1. We revealed the wide prevalence of CpDV, mostly at low densities, in *Cx. pipiens* laboratory lines.
2. We monitored CpDV presence in several cell types in ovaries including the oocytes where they co-exist with the bacterial endosymbiont *Wolbachia*.
3. We demonstrated that CpDV density in ovaries was influenced by *Wolbachia* presence and strain variations
4. We studied CpDV vertical transmission depending on *Wolbachia* presence and strain variations

We report for the first time the occurrence of widely distributed covert CpDV infections in *Cx. pipiens* laboratory colonies. CpDV is maintained by both vertical and horizontal transmission and under the influence of the vertically transmitted bacterial endosymbiont *Wolbachia.* Taken together our results suggest that CpDV could be very prevalent in nature putatively influencing natural *Cx. pipiens* populations’ dynamics.

**SUMMARY:** *Culex pipiens* densovirus (CpDV), a single stranded DNA virus, has been isolated from *Culex pipiens* mosquitoes but differs from other mosquito densoviruses in terms of genome structure and sequence identity. Its transmission from host to host, the nature of its interactions with both its host and host’s endosymbiotic bacteria *Wolbachia* are not known. Here we report the presence of CpDV in the ovaries and eggs of *Cx. pipiens* mosquitoes in close encounters with *Wolbachia*. In the ovaries, CpDV quantity significantly differed between mosquito lines harboring different strains of *Wolbachia* and these differences were not linked to variations in *Wolbachia* densities. CpDV was vertically transmitted in all laboratory lines to 17%-20% of the offspring. For some females, however, the vertical transmission reached 90%. Antibiotic treatment that cured the host from *Wolbachia* also significantly affected both CpDV quantity and vertical transmission suggesting an impact of host microbiota, including *Wolbachia*, on CpDV transmission. Overall our results show that CpDV is transmitted vertically *via* transovarian path along with *Wolbachia* with which it shares the same cells. Our results are primordial to understand the dynamics of densovirus infection, their persistence and spread in populations considering their potential use in the regulation of mosquito vector populations.

## Introduction

Densoviruses are non-enveloped, single stranded DNA viruses. They constitute a diverse subfamily within the *Parvoviridae* family that includes both vertebrate (*Parvovirinae*) and invertebrate (*Densovirinae*) viruses (Cotmore *et al.*, 2014; Tijssen *et al.*, 2016). Isolated from many insects (e.g. butterflies, moths, mosquitoes, crickets, grasshoppers and cockroaches) and crustaceans (e.g. prawns and crayfish), they are thought to be widely distributed among arthropods (Kerr *et al.*, 2005; Tijssen *et al.*, 2016). Recent studies, investigating publicly available transcriptomic and genomic databases, have shown their presence in even broader range of non-vertebrate hosts (e.g., molluscs, annelids, nematodes, cnidarians and sea stars)(François *et al.*, 2016; Kang *et al.*, 2017). In arthropods, most of the densoviruses known so far exhibit a tissue polytropism replicating in many tissues-such as the fat body, hypodermis, epidermis, central nervous system, muscular and tracheal cells, gut, hemocytes, ovaries- and are considered to be lethal for their natural hosts (Tijssen *et al.*, 2016). Due to their observed pathogenicity, narrow host range and sustainability out of their host-cells in nature, they attracted scientific attention as biological control tools especially against mosquitoes (Carlson *et al.*, 2006; Mutuel *et al.*, 2010; Gosselin Grenet *et al.*, 2015; Suzuki *et al.*, 2015; Johnson and Rasgon, 2018).

Mosquito densoviruses (MDVs) have been described in natural populations (Afanasiev *et al.*, 1991; Boublik *et al.*, 1994; Kittayapong *et al.*, 1999; Rwegoshora *et al.*, 2000; Ng *et al.*, 2011), laboratory colonies (Jousset *et al.*, 2000) and established mosquito cell lines (Jousset *et al.*, 1993; O’Neill *et al.*, 1995). The first described mosquito densoviruses were *Aedes aegypti* densovirus (AeDV) (Lebedeva *et al.*, 1972) and *Aedes albopictus* densovirus (AalDV) (Jousset *et al.*, 1993). Thereafter many other MDV strains, namely *Toxorhynchites amboinensis* densovirus (TaDV), *Haemagogus equinus* densovirus (HeDV), *Ae. aegypti Thai strain* densovirus (AThDV), *Ae. Peruvian densovirus* (APeDV) have been identified (O’Neill *et al.*, 1995; Kittayapong *et al.*, 1999; Rwegoshora *et al.*, 2000; Ledermann *et al.*, 2004). Studies on MDV pathogenicity and transmission modes have generally shown a dose and age dependent infectivity and virulence (Barreau *et al.*, 1996; Kittayapong *et al.*, 1999; Ledermann *et al.*, 2004; Rwegoshora and Kittayapong, 2004; Becnel and White, 2007). Vertical transmission of these viruses has only been observed when using sub-lethal doses that reduced the adult mosquitoes’ longevity and fertility but did not kill them (O’Neill *et al.*, 1995; Barreau *et al.*, 1996, 1997; Kittayapong *et al.*, 1999; Rwegoshora and Kittayapong, 2004; Becnel and White, 2007). In contrast, *Anopheles gambiae densovirus* (AgDV), a recently discovered MDV, which exhibits a negligible pathogenic effect on its host *A. gambiae* mosquitoes (Ren *et al.*, 2008, 2014) has been shown to replicate in its host and to be vertically transmitted to the offspring highlighting its potential as transducing agents for mosquito control (Ren *et al.*, 2008; Suzuki *et al.*, 2014).

All of the abovementioned MDVs described so far are closely related (82-99% sequence homology), and are considered as strains from two distinct species (dipteran brevidensovirus 1 and 2) belonging to the genus *Brevidensovirus* (Cotmore *et al.*, 2014). To date, the only known densovirus to infect mosquitoes, but does not belong to *Brevidensovirus* genus, is the *Culex pipiens* densovirus (CpDV). CpDV is a dipteran ambidensovirus (Jousset *et al.*, 2000; Baquerizo-Audiot *et al.*, 2009) which differs significantly from dipteran brevidensoviruses by the lack of sequence homology (<30% in Non-structural (NS1/Rep) gene) and antigenic cross-reactivity (Jousset *et al.*, 2000; Baquerizo-Audiot *et al.*, 2009). It belongs to *Ambidensovirus* genus along with lepidopteran densoviruses, such as *Junonia coenia* densovirus (JcDV) and *Galleria melonella* densovirus (GmDV). Ambidensoviruses have not yet been described in other dipteran species suggesting that they could have been acquired in *Cx. pipiens* by horizontal transfer. CpDV indeed shares the characteristics of other ambidensoviruses having a larger genome size than brevidensovirus (~6kb compared to ~4kb), an ambisense genome organization and significant sequence homology at both capsid and NS gene levels (Jousset *et al.*, 2000; Baquerizo-Audiot *et al.*, 2009).

Two decades after its first isolation from *Cx. pipiens* laboratory colonies, due to an unusually high larval mortality, we detected the presence and persistence of CpDV in all life stages of seemingly healthy *Cx. pipiens* lines by a PCR diagnostic test (data not shown). This long-term persistence of CpDV in laboratory colonies through generations certainly relies on an efficient transmission. Transmission modes are generally categorized as i) horizontal, transmission from host to host via the environment, ii) vertical transmission from the host parent to offspring, iii) mixed-mode, both horizontal and vertical transmission (Ebert, 2013). Transmission modes have long been investigated in many symbiotic systems especially to understand their impact on virulence evolution (Ewald, 1983, 1987; Alizon *et al.*, 2009; Cressler *et al.*, 2016; Le Clec’h *et al.*, 2017). Empirical studies mostly showed that increased horizontal transmission led to increased virulence, while dominance of vertical transmission mode mostly reduced the virulence [reviewed in (Cressler *et al.*, 2016)]. Therefore transmission modes can have a major impact on the dynamics of virus infection, their persistence and spread in populations.

To understand CpDV infection dynamics, especially its maintenance through generations, we investigated the ability of *Cx. pipiens* mosquitoes to transmit CpDV vertically. *Cx. pipiens* are all naturally infected with *Wolbachia* strains belonging to one of the five genetically distinct phylogenetic group wPipI to wPipIV (Atyame et al. 2011). These *wPip* strains induce a wide diversity of cytoplasmic incompatibility phenotypes in their mosquito host (Atyame et al. 2014). In addition to their ability to manipulate host reproduction to benefit their own vertical transmission (Stouthamer *et al.*, 1999; Werren *et al.*, 2008; Sicard *et al.*, 2014), *Wolbachia* are also known to interact with several of their hosts’ viruses (Hedges *et al.*, 2008; Teixeira *et al.*, 2008; Zug and Hammerstein, 2014). While putative difference in protective ability between the *w*Pip groups identified in *Cx. pipiens* has not yet been investigated, a general protection by wPip has been shown against West Nile Virus infection (Micieli and Glaser, 2014). To date, no *Wolbachia-mediated* “protection” against DNA viruses have been shown in any insects. Contrarily, one study demonstrate that *Wolbachia* makes their lepidopteran host more susceptible against a Baculovirus (Graham *et al.*, 2012).

We thus examined the influence of *Wolbachia* on CpDV density and vertical transmission. To do that, we monitored CpDV density and location in the ovaries and eggs, and then quantified its vertical transmission in *Cx. pipiens* mosquitoes naturally infected with both *Wolbachia* and CpDV. Given their importance in *Cx. pipiens*’ phenotype both in natural populations and laboratory colonies, we took *Wolbachia* into account in each step of the study and further investigated the possible effects of their absence on CpDV presence and vertical transmission.

## Results

### CpDV presence in ovaries and eggs

CpDV was always present, along with *Wolbachia*, in the ovaries of females from ten laboratory lines tested by qPCR and its amount varied between different lines (glm, df=13, dev=82.448, p<0.001, Fig. 1). Only two pools of ovaries-one from Ich13 and another from Ich09 line-over 50 pools tested were CpDV negative. We did not detect any statistically significant effect of *Wolbachia* amount (glm, df=1, dev=0.007, p= 0.967, Fig. 1A) on the amount of CpDV in ovaries. To create backcrossed lines for further experimentation, we used “Harash” mosquito line with the highest amount of CpDV in ovaries (120±114 CpDV/host genome, Supplementary Table 1, Fig. 1B) and “Tunis” line with less CpDV in their ovaries (0.005± 0.002 CpDV/host genome, Tunis, Supplementary Table 1, Fig. 1B).

**Figure 1.**
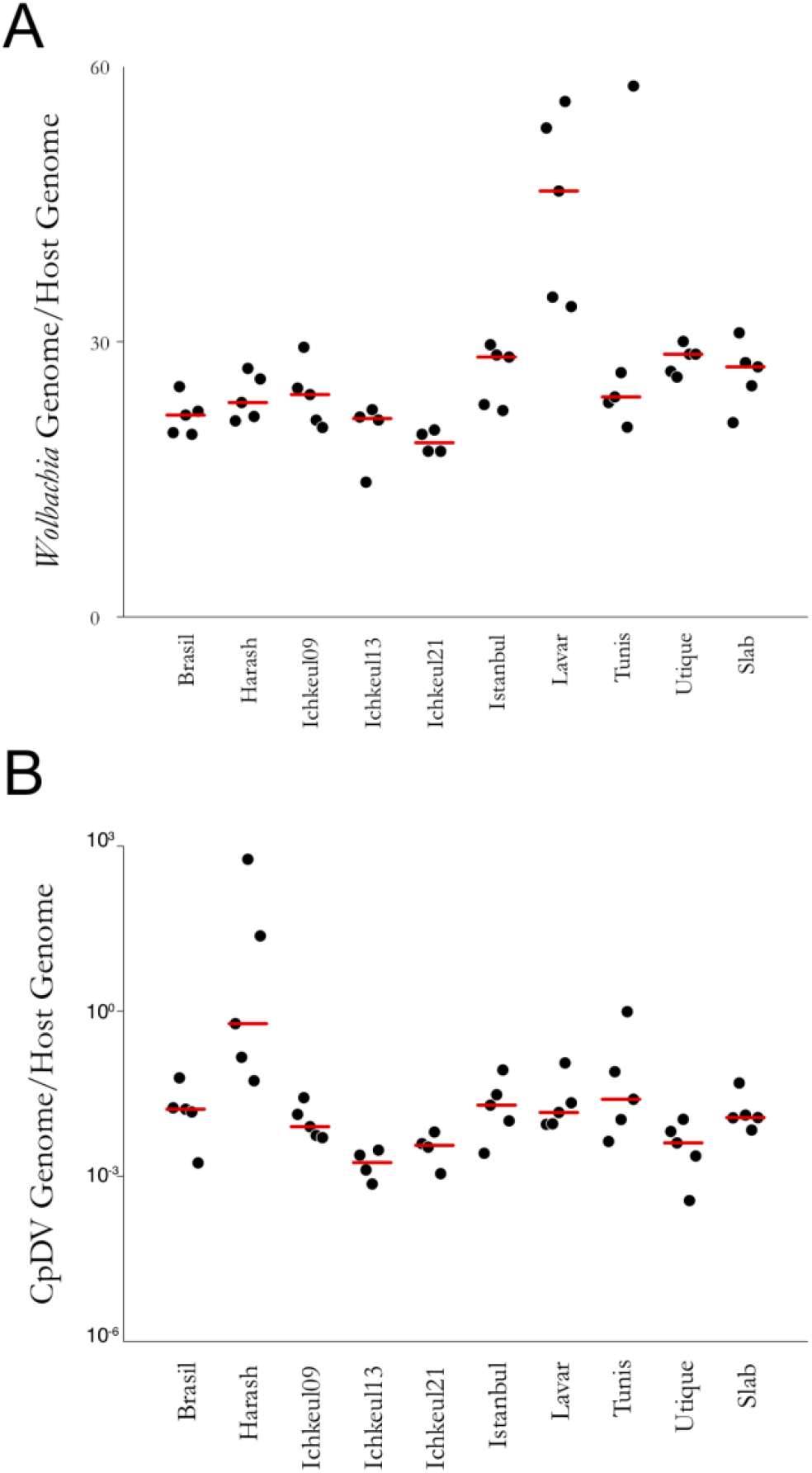
The two symbionts *Wolbachia* and CpDV coexist in *Cx. pipiens* ovaries of ten laboratory lines *Wolbachia* amount. **(A)** Ovaries from three females per line were dissected and pooled. *wsp* copy number/*ace-2* copy number was quantified for 5 pools per line by qPCR assays. **CpDV amount (B)** NS2 copy number and *ace-2* copy number have also been quantified in the same pools. Harash line with the highest quantity of CpDV in ovaries and Tunis line with less CpDV in ovaries were selected for further experimentation.

To confirm the presence of virus particles -and not only of the viral genome (*i.e*. as detected by qPCR)- and to investigate their localization in the ovaries, we dissected ovaries from 6 days old Harash females and performed immunolabelling of CpDV using an anti-capsid antibody for confocal observation (Fig. 2 and Supplementary Fig. 1). The labelling of CpDV was mainly observed in follicular cells surrounding the follicles (Fig. 2A-B-C) and occasionally in some muscle cells of the sheath enclosing the ovariole (Fig. 2D). In both cell types, the labelling is mainly nuclear with some dots present in the cytoplasm of follicular cells (Fig. 2B to E). Within the follicles, the labelling of CpDV in the nurse cells was not as apparent and rather similar to the background immunofluorescence (Fig. 2D-E). Although we cannot entirely exclude a lack of permeability and accessibility of the nurse cells to the antibodies, a clear labelling was nevertheless observed in a well-defined area of the nucleus of the oocyte (Fig. 2E). As all the *Cx. pipiens* lines were infected, the specificity of the CpDV labeling was tested on ovaries isolated from PCR tested CpDV-free *Ae. albopictus* mosquitoes and no labelling was detected (Supplementary figure 2).

**Figure 2.**
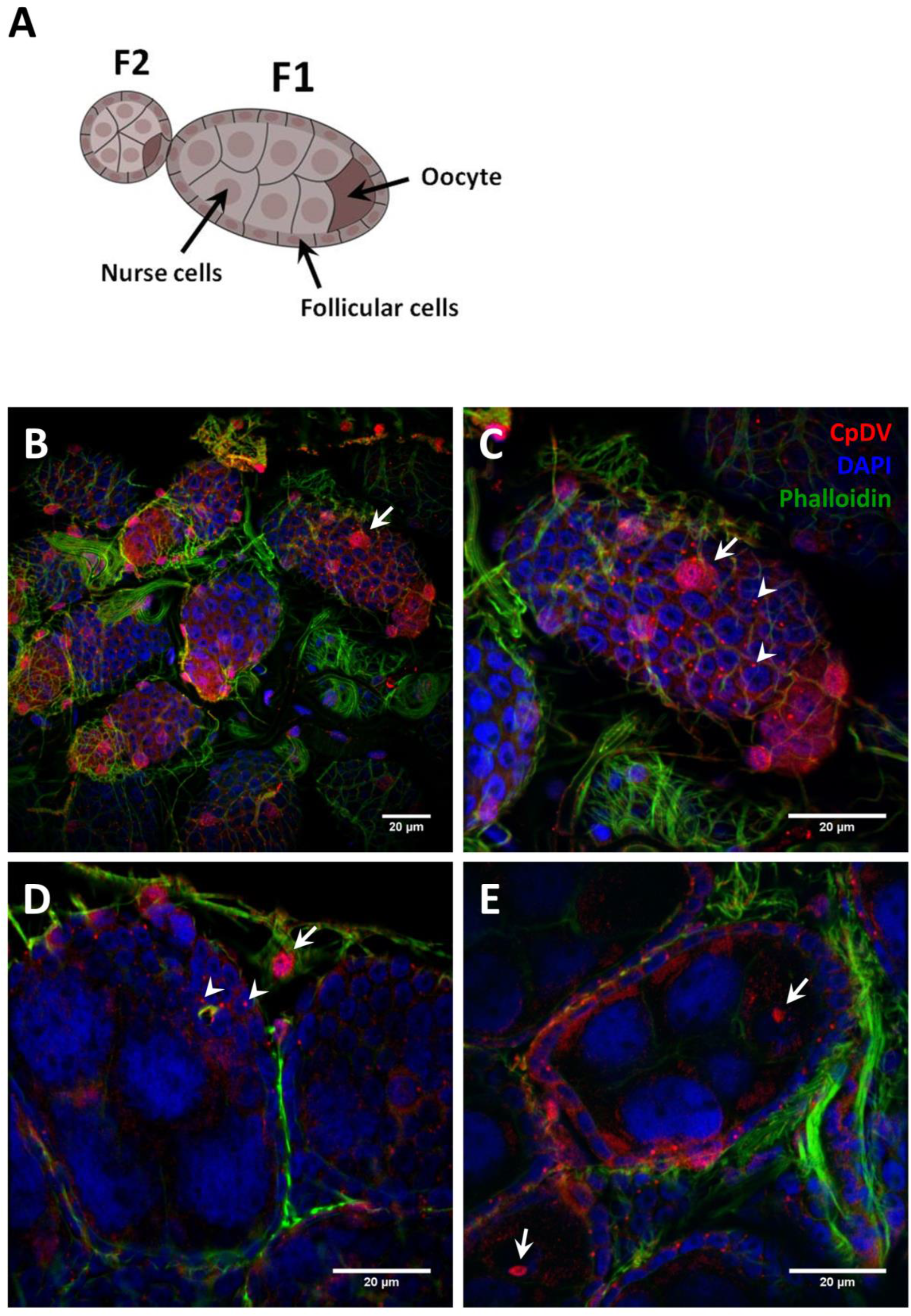
Immunolabeling of CpDV in *Cx. pipiens* ovaries. **Structure of a *Cx. pipiens* ovariole (A).** The mosquito ovaries are paired and composed of functional units called ovarioles, containing primary (F1) and secondary follicles (F2) (Clements, 1992). Each primary follicle contains seven nurse cells and one oocyte, both surrounded by follicular epithelium. The secondary follicle is composed of undifferentiated cells surrounded by follicular epithelium. The secondary follicle will turn into a primary follicle after oviposition of the oocyte contained in the previous primary follicle (Raikhel and Lea, 1983). The sheath that enclosed each ovariole is a thin membrane with an outer muscle layer composed of isolated cell bodies interconnected by muscle fibers. **Confocal images of ovaries isolated from 6 days old *Cx. pipiens* females (B,C,D,E).** Immunolabeling of CpDV (in red) was performed with an anti-capsid antibody, actin was labeled with phalloidin-FITC (green) and nuclei with DAPI (blue). Arrows show CpDV-infected cells and arrowheads point cytoplasmic dots of CpDV labelling. Images of the surface of an ovary and an ovariole (composed of a primary and a secondary follicles) are shown in B and C, respectively. Internal views of primary follicles are shown in D and E.

To examine the CpDV replication, which takes place in the nucleus, we prepared ultrathin sections of ovaries and analyzed the nuclear ultrastructure with transmission electron microscopy (TEM) (Fig. 3). Although no unequivocal densonucleosis was observed, follicular and nurse cells showed hypertrophied nuclei (Fig. 3A and G). Intranuclear structures that could be interpreted as viral factories were observed in several follicular cells (Fig. 3B), nurse cells (Fig. 3E) and oocytes (Fig. 3H). Particles with the expected diameter of CpDV (about 25 nm) were also present in the nucleoplasm (Fig. 3C and F). To confirm the presence of viral particles in ovaries, we prepared filtered homogenate of the full organ for observation using TEM. Micrographs revealed the presence of both icosahedral empty capsids and CpDV virions with the expected diameter (Fig. 3I).

**Figure 3.**
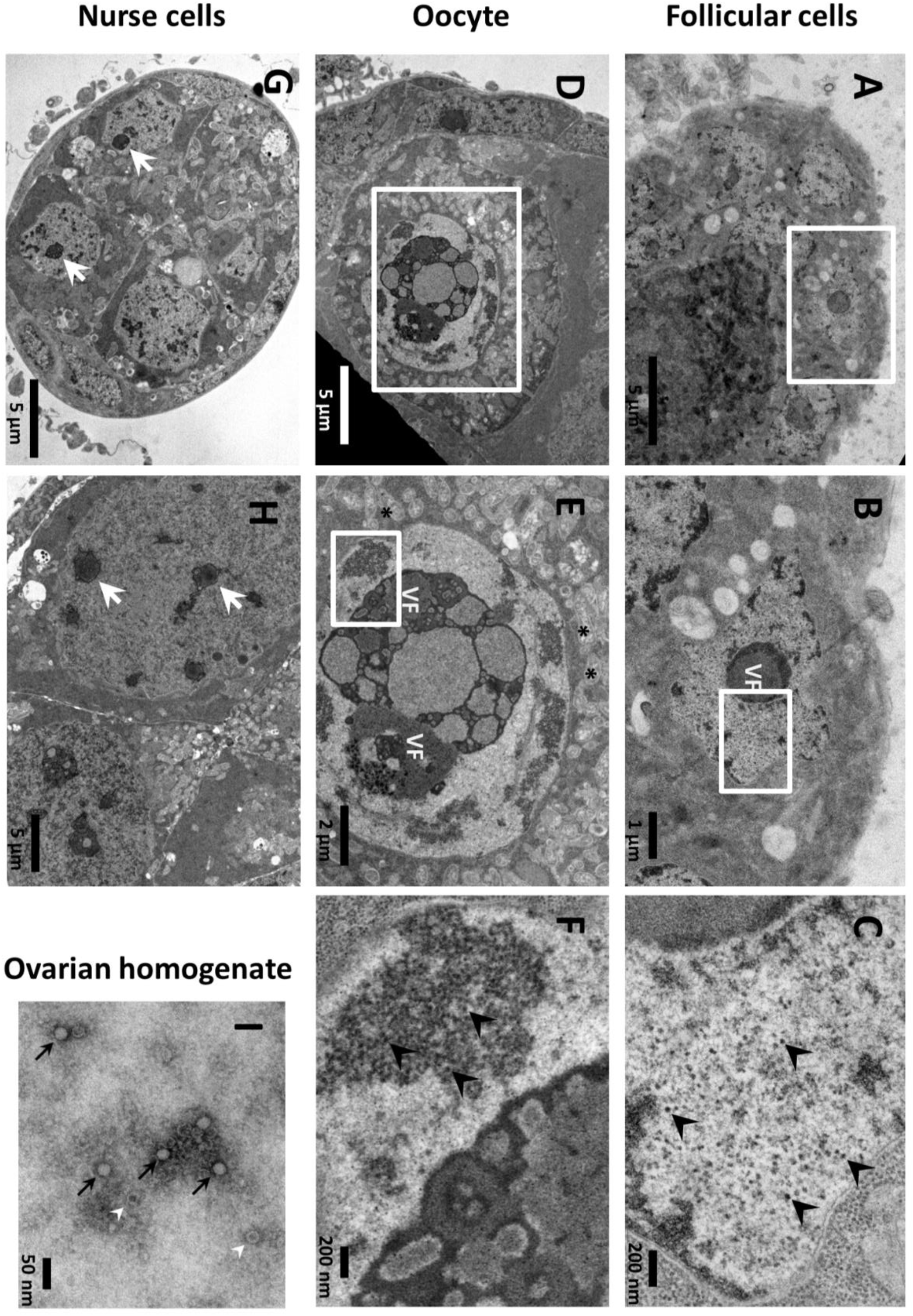
Electron micrographs of *Cx. pipiens* ovaries. **Ultrathin sections of ovaries isolated from 6 days old females (Harash line) (A to H).** Large views of an ovariole (primary follicle) in A, D and G (bar scale: 5 μm). Ultrastructure of the hypertrophied nuclei and putative virus factories in a follicular cell (A and B), in an oocyte (D and E) and in a nurse cell (G and H). Zooms in the nucleoplasm of the follicular cell **(C)** and in the oocyte (F) show particles with the expected size of CpDV (25 nm). **Virus particles observed in a filtered homogenate of ovaries (I).** Arrows show icosahedral virus particles and arrowheads point empty particles with the expected size of CpDV (bar scale: 50 nm).

Same experiments were performed on freshly laid dechorionated eggs from Harash females (Fig. 4). Immunolocalization of CpDV and DAPI staining of *Wolbachia* cells at early stage of the embryo development showed a *Wolbachia* localization especially in the poles, and a CpDV localization both inside and in the periphery of the eggs suggesting no co-localization of these two (Fig. 4A, Top Panel). At the syncytial blastoderm stage, CpDV were co-localized with the nuclei, presumably starting viral replication (Fig. 4A, Bottom Panel). *Wolbachia* were easily detected in the cytoplasm in the ultrathin sections of fertilized eggs even at an early stage of development (Fig. 4C). However, at the early stage of development nuclei were rare, meaning CpDV could only be observed in the cytoplasm where the ribosomes are also present (Fig. 4C). As ribosomes and CpDV have the same diameter, it is not possible to firmly distinguish them (Fig. 4C).

**Figure 4.**
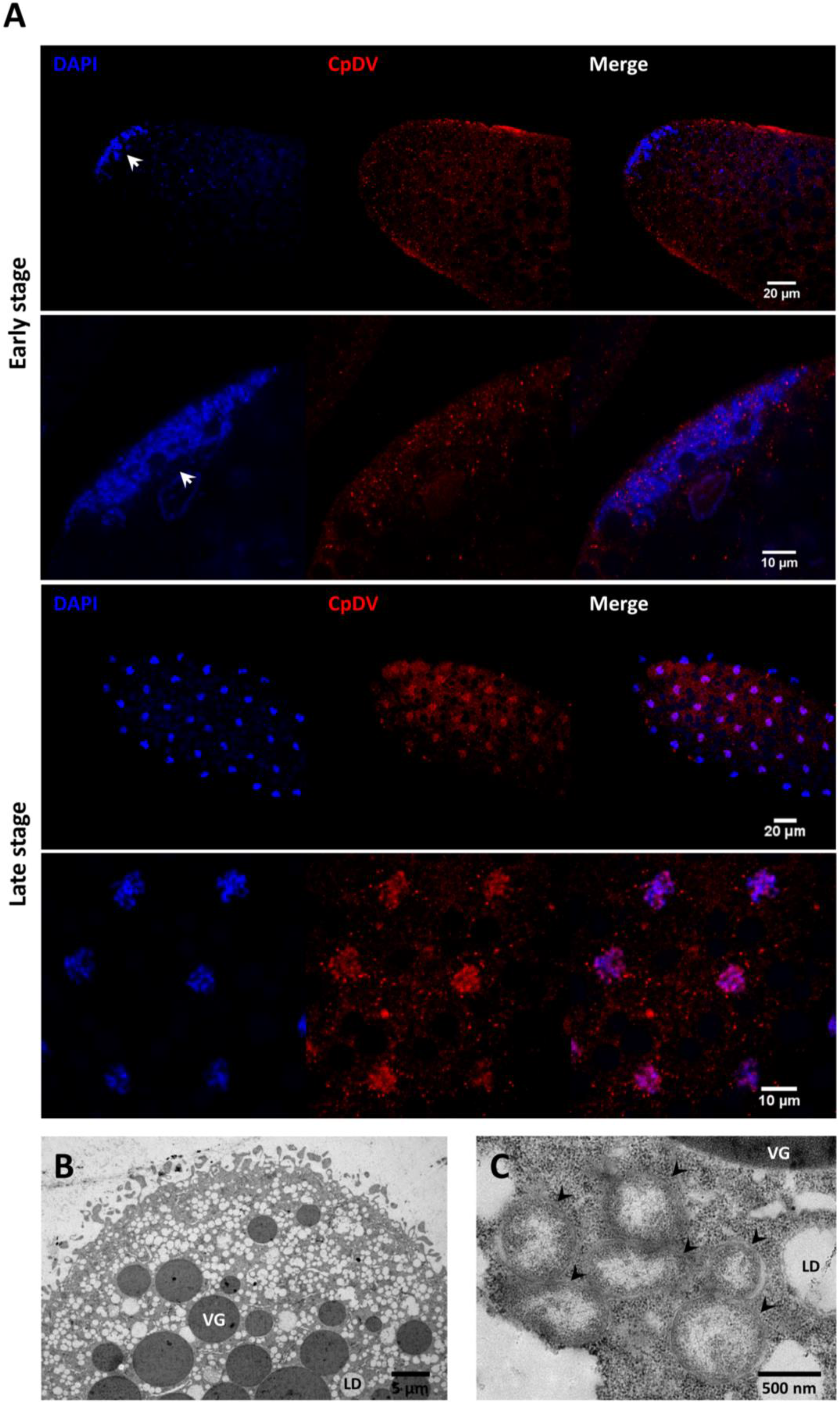
Localization of CpDV in *Cx. pipiens* eggs. **Confocal images of dechorionated eggs from *Cx. pipiens* females (Harash line) (A).** CpDV (in red) were immunolabeled with an anti-capsid antibody; *Wolbachia* and nuclei (in blue) were labeled with DAPI. Top panel shows eggs in the first hour after laying (early developmental stage) and bottom panel eggs 2 hours after laying (late developmental stage). The second part of the Top panel is a zoom of one egg pole in the first hour after laying; the second part of the Bottom panel is a zoom in the egg cytoplam. Arrows show *Wolbachia* cells at one egg pole. **Electron micrographs of ultrathin sections of dechorionated eggs (B and C).** In B, a large view of an early egg pole (bar scale: 5 μm), showing the accumulated reserves required for the embryo development (VG, vitellin granule; LD: lipid droplet). In C, a zoom (bar scacle: 500 nm) inside the cytoplasm of the early egg showing numerous *Wolbachia* (arrowheads). The virus particles cannot be distinguished from the ribosomes because of their similar size.

### Effect of infection with distinct *Wolbachia* strains on CpDV amount in ovaries

To assess the effect of *Wolbachia* (strains and density) on the amount on CpDV in the ovaries, we have established mosquito lines, with help of a backcrossing protocol, with the same host nuclear genetic background but different *Wolbachia* strains (BcTunis, Slab genetic background harbouring *w*Pip I from Tunis line; BcHarash, Slab genetic background harbouring wPip IV from Harash line, Table 1). We then quantified the amount of *Wolbachia* and CpDV in these lines as well as in Harash, Tunis and Slab lines (Fig. 5). *Wolbachia* amount in the ovaries was not significantly different between the *Cx. pipiens* lines no matter what the *Wolbachia* strain and host genetic background were (*Wolbachia* strain: glm, df=2, dev=17.703, p=0.875; host genetic background: df=1 dev=0.034, p=0.981, Fig. 5A). Indeed, there was no significant effect of the *Wolbachia* amount in the ovaries on CpDV density (lme, dev=0.001, df=1, p=0.9). However, *Wolbachia* strain (*i.e*. the genotype) infecting the mosquito lines explained a significant part of the variance in the CpDV amount in the ovaries (lme, dev=12.236, df=1, p<0.001, Fig. 5B) while the host genetic background had no significant effect on CpDV amount in the ovaries (lme, dev=1.602, df=2, p= 0.449; Fig. 5B).

**Figure 5.**
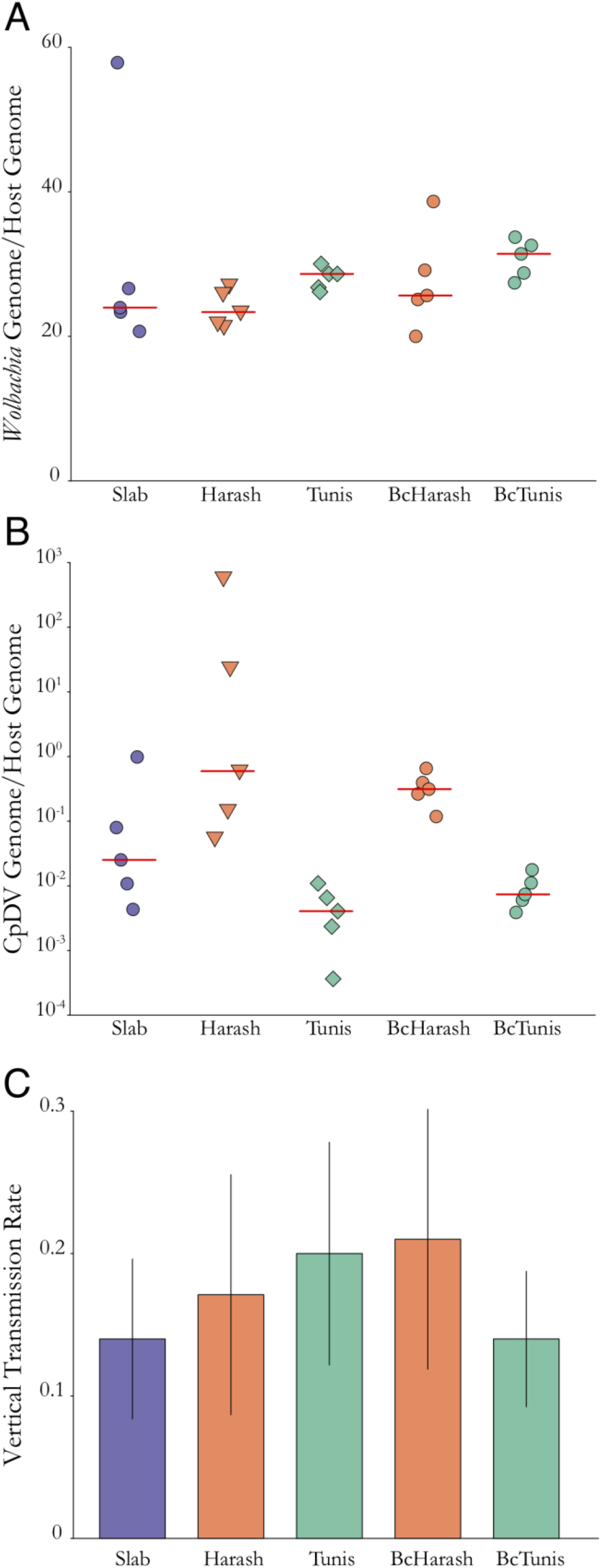
*Wolbachia* and CpDV amounts in the ovaries and vertical transmission of CpDV in different *Cx. pipiens* lines with similar genetic background but different *Wolbachia* strains. Slab line’s genetic background was introgressed into Harash and Tunis lines, to establish BcHarash and BcTunis lines with the same genetic background as Slab (shown in circles) and same *Wolbachia* strain as Harash (shown in orange) or Tunis (shown in green) respectively. ***Wolbachia* amount in ovaries (A).** *Wolbachia* amount in the ovaries was not significantly different between the *Cx. pipiens* lines no matter what the *Wolbachia* strain and host genetic background were (*Wolbachia* strain: glm, df=2, dev=17.703, p=0.875; host genetic background: df=1 dev=0.034, p=0.981). **CpDV amount in ovaries (B)** Ovaries from three females were dissected and pooled. 5 pools per line have been tested by qPCR assays to quantify CpDV and *Wolbachia* amount. Mixed effect linear model showed no effect of *Wolbachia* amount or genetic background on the CpDV amount in the ovaries (*Wolbachia* amount dev=0.001, df=1, p=0.9, genetic background: dev=1.602, df=2, p= 0.449). *Wolbachia* strain, however significantly affected the CpDV amount (dev=12.236, df=1, p<0.001). **CpDV vertical transmission rate (C)** Females from each line were collected 5 days after their blood meal and isolated to lay eggs. Resulting L1 offspring was tested for CpDV prevalence and vertical transmission rate was calculated per female. A binomial generalized mixed effect model was fitted with host genetic background, *Wolbachia* strain as well as CpDV and *Wolbachia* amount in females’ bodies as fixed effects and replicates as random effects. None of these variable had a significant effect on the vertical transmission rate.

### Effect of infection with distinct *Wolbachia* strains on CpDV vertical transmission

Using the same mosquito lines, we tested the vertical transmission of CpDV. For this, we let 10 females per line to lay eggs in isolation. To estimate a vertical transmission rate per female mosquito, both females and 10 L1 stage larvae per female were collected and tested for CpDV presence. All the mothers were CpDV positive. CpDV were transmitted to an average of 17% of the offspring (n=49, 0.172 ± 0.0317, Fig5C). However, the vertical transmission rate highly diverged between individuals ranging from 0% (n=19 out of total 49 checked) to 90% (n=1, out of total 49 checked). This inter-individual variability was not significantly explained neither by the host genetic background (df=2, dev=0.278, p= 0.870, Fig5C), *Wolbachia* strain (df=1, dev=0.152, p= 0.696, Fig3C), *Wolbachia* amount (df=1, dev= −0.135, p=0.713, Fig5C) nor by CpDV amount in the whole body of the parental females (df=1, dev=-0.180, p=0.672, Fig5C).

### Effect of *Wolbachia* curing by tetracycline on the amount of CpDV in ovaries

As all the *Cx. pipiens* isofemale lines were stably infected with wPip (Wo+), to test the effect of the *Wolbachia* absence on CpDV presence and vertical transmission, we treated our Slab, BcHarash, and BcTunis lines, all having the same genetic background, with tetracycline (TC) for three generations to cure *Wolbachia*. We quantified *Wolbachia* and CpDV in dissected ovaries and carcasses of Wo+ (before TC treatment) and Wo- females (after TC treatment), as well as females during the 1^st^ and 3^rd^ generation of TC treatment (Fig. 6). There was no significant change between 1^st^ and 3^rd^ generations in terms of CpDV amount (t = −0.643, df = 25.918, p-value = 0.525, Fig. 6B). We thus grouped them together in a “TC treatment” group for further statistical analyses and figures.

**Figure 6.**
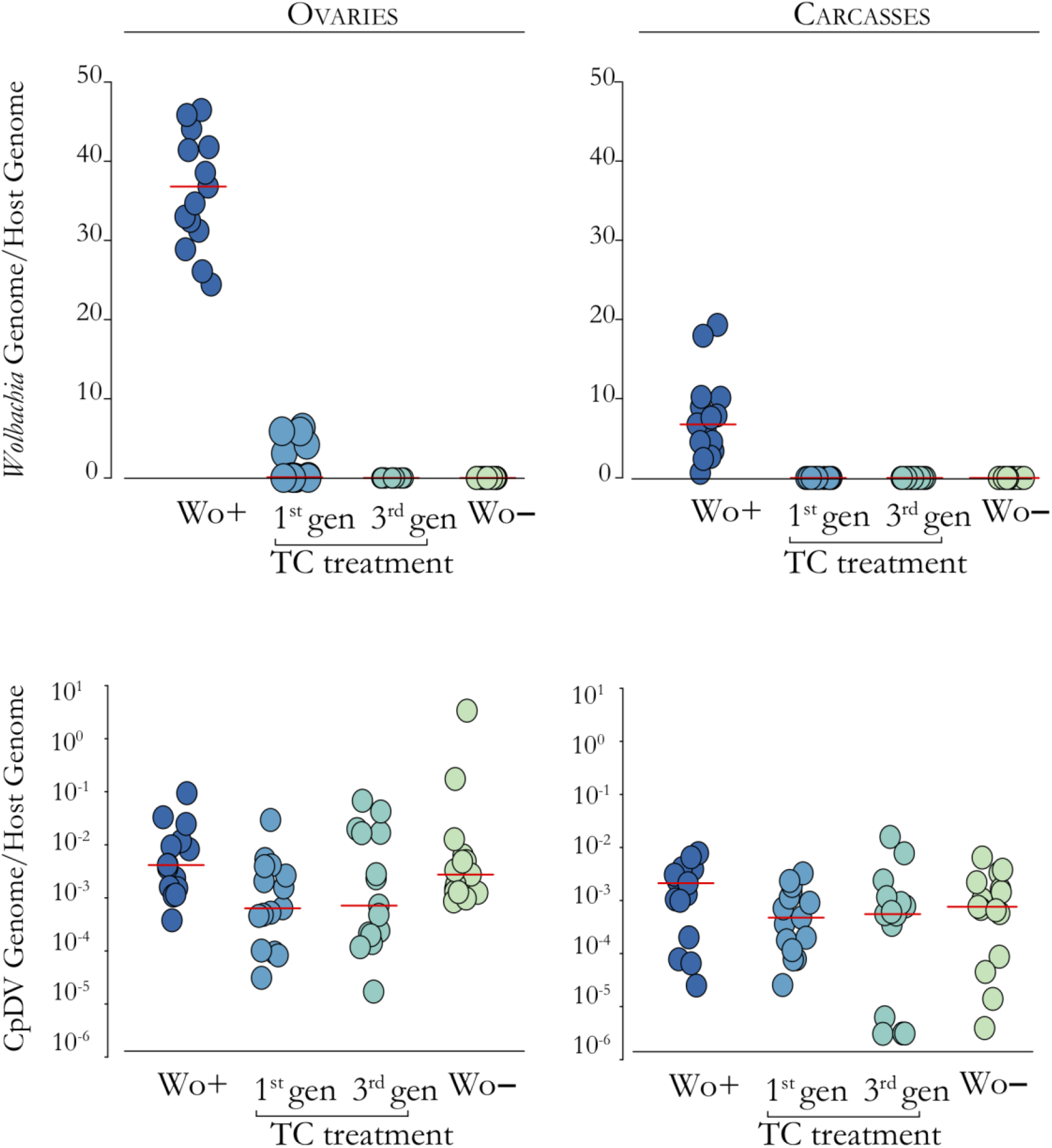
Effect of curing *Wolbachia* by tetracycline treatment on CpDV amount in *Cx. pipiens*. We have treated Slab, BcHarash and BcTunis lines with tetracycline (TC) for three generations and dissected the ovaries of 6 days old females. Dissections were performed on the non-treated mosquitoes (Wo+), first and third generation of mosquitoes during tetracycline treatment (during TC treatment) and three generations after the tetracycline treatment was finished (Wo-). **Effect of tetracycline treatment on the amount of *Wolbachia* in ovaries and carcasses (A).** *Wolbachia* amount decreased immediately starting from first generation during tetracycline treatment. Even though *Wolbachia* was detected in the ovaries of some mosquitoes from the first generation of the treatment, there were no *Wolbachia* detected in the subsequent treatment generation neither in ovaries nor in carcasses. **Effect of tetracycline treatment, leading to full *Wolbachia* disappearance, on CpDV amount (B).** CpDV amount in the ovaries significantly decreased starting from the first round of tetracycline treatment but increased when tetracycline treatment stopped and the line “recovered” from the treatment (Wo-)(Dunnett’s test with Bonferroni Correction; Wo+ vs. during TC treatment, t=2.302, p=0.025; Wo+ vs. Wo-, t= 0.118, p=1). Carcasses had less CpDV compared to ovaries (Wilcoxon rank sum test, W = 168, p = 0.021). The effect of the treatment was not significant on the carcasses (Wilcoxon rank sum test with Bonferroni correction; Wo+ vs. during TC treatment, W= 132.5, p=0.08; Wo+ vs. Wo-, W= 100, p=0.138; during TC treatment vs. Wo-, W=308, p=1).

Prior to tetracycline treatment *Wolbachia* were found in higher density in the ovaries compared to the carcasses (Wilcoxon rank sum test, W= 225, p<0.001; Fig. 6A). By the end of the third generation of tetracycline treatment, as expected, *Wolbachia* totally disappeared both from the ovaries and the carcasses (Fig. 6A).

In non-treated Wo+ females, higher amount of CpDV were measured in ovaries compared to carcasses (Wilcoxon rank sum test, W = 168, p = 0.021, Fig. 6B). In the carcasses, TC treatment did not cause any significant effects on CpDV amount (Fig. 6B, Wilcoxon rank sum test with Bonferroni correction; Wo+ vs. during TC treatment, W= 132.5, p=0.08; Wo+ vs. Wo-, W= 100, p=0.138; during TC treatment vs. Wo-, W=308, p=1). On the contrary, in the ovaries CpDV amount significantly decreased during the TC treatment (Dunnett’s test with Bonferroni Correction; Wo+ vs. during TC treatment, t=2.302, p=0.025; Fig. 6B). However, in Wo- lines (three generations after the end of the tetracycline treatment) CpDV amount increased and reached a level not significantly different from non-treated Wo+ lines (Dunnett’s test with Bonferroni Correction; Wo+ vs. Wo-, t= 0.118, p=1).

### Effect of *Wolbachia* curing by tetracycline on CpDV vertical transmission rate

CpDV vertical transmission decreased significantly from an average of 25% to an average of 13% during tetracycline treatment (Wilcoxon rank sum test with Bonferroni correction; Wo+ vs. TC treatment, W= 488, p= 0.004, Fig. 7). Three generations after the end of tetracycline treatment, vertical transmission in Wo- lines stayed marginally lower than non-treated Wo+ lines (Wilcoxon rank sum test with Bonferroni correction; Wo+ vs. Wo-, W= 576.5, p= 0.05, Fig. 7).

**Figure 7.**
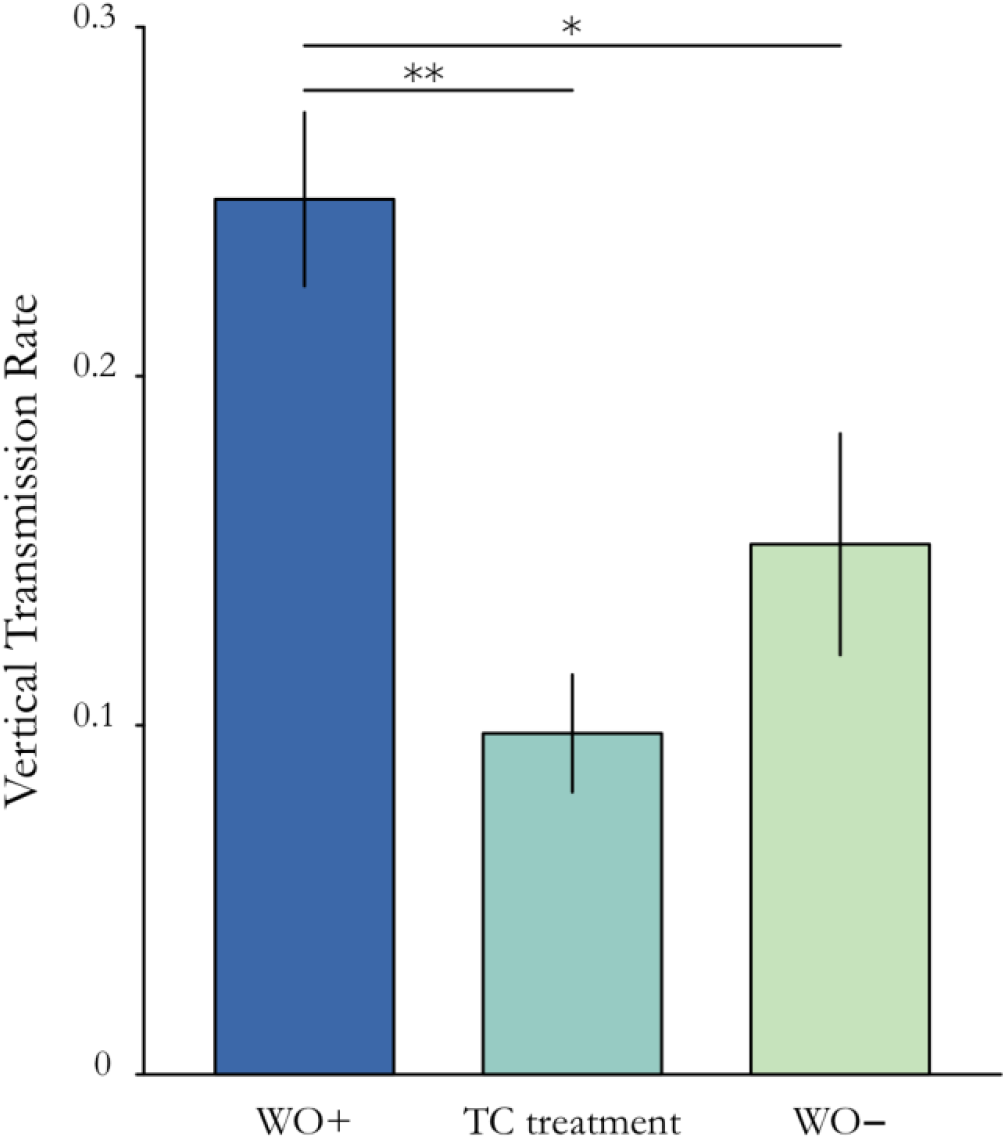
Effect of curing *Wolbachia* by tetracycline treatment on CpDV vertical transmission. In BcHarash, BcTunis and Slab lines, Females emerged from the larvae that underwent the first tetracycline treatment were isolated five days after bloodmeal to lay eggs. 10 females per line (total n=30), 10 L1 stage larvae per female were collected and tested for CpDV presence to estimate vertical transmission rate. This procedure was repeated for females emerged after the third generation of TC treatment and for the females Wo- (three generation after the end of tetracycline treatment). There was a significant decrease in vertical transmission during tetracycline treatment (Wilcoxon rank sum test with Bonferroni correction; Wo+ *vs*. during TC treatment, w= 488, p= 0.004). Three generations after end of the treatment the vertical transmission rate increased but still was marginally significant from the non-treated group (Wilcoxon rank sum test with Bonferroni correction; Wo+ vs. Wo-, w= 576.5, p= 0.05).

## Discussion

CpDV was isolated for the first time from *Cx. pipiens* larvae almost two decades ago (Jousset *et al.*, 2000). The features of densonucleosis (hypertropied nuclei and electron-dense virogenic stroma) was then observed in larval tissues (hypodermis, fat body, midgut, imaginal disks and nervous tissue) suggesting CpDV’s role on the larval mortality (Jousset *et al.*, 2000). Here, we showed that CpDV was also present in apparently healthy adults especially in their ovaries along with endosymbiotic bacteria *Wolbachia*. Even if the densonucleosis was not unequivocal, intranuclear structures similar to viral factories were observed and viral particles were isolated from ovaries. Furthermore immunolabelling revealed virus particles in the follicular cells and the oocytes. Only couple of studies reported the presence of densoviruses in the ovaries. *Helicoverpa armigera* densovirus (*Ambidensovirus*) and *A. gambiae* densovirus (*Brevidensovirus*) have been shown to reach high densities in ovaries of their hosts while they were also able to infect other host tissues (Ren *et al.*, 2008; Suzuki *et al.*, 2014; Xu *et al.*, 2014).

The presence of CpDV capsids in the ovaries and eggs highlighted the possibility of vertical transovarial transmission. Transmission mechanisms of densoviruses are generally thought to be horizontal causing epizootics (Tijssen *et al.*, 2016). Nevertheless, vertical transmission has been shown both for some ambidensoviruses and brevidensoviruses. Interestingly, ambidensoviruses with different vertical transmission rates had different types of symbiotic interactions: while the 100% vertically transmitted ambidensovirus of *H. armigera* is mutualistic (Xu *et al.*, 2014), the *Myzus persicae* densovirus which is transmitted to 30-40% of the offspring has been shown to affect its hosts adversely (Van Munster *et al.*, 2003). In mosquitoes experimentally infected with brevidensoviruses, the vertical transmission ranged between 28 to 62%: i) for *Ae. albopictus* densovirus, the vertical transmission rate ranged between 28-55% depending on the virus titer in parental females (Barreau *et al.*, 1997); ii) for *Ae. aegypti* densovirus vertical transmission ranged from 42% to 62% depending on the timing of the horizontal infection of the larvae (the larvae that were infected during L3 stage transmitted the virus less, as they became adults, than the larvae infected during L1 stage) (Suchman *et al.*, 2006), and iii) for *A. gambiae* densovirus vertical transmission was detected in 28% of larval offspring (Ren *et al.*, 2008). We showed that CpDV was transmitted to an average of 17% of the larval offspring in naturally infected *Cx. pipiens* lines. Even if the vertical transmission rate varied between individuals, this rate was quite stable (15%-21%) between the *Cx. pipiens-Wolbachia* lines tested through the different replicates. The importance of inter-individual variability in vertical transmission rate of CpDV revealed the complexity of the factors affecting vertical transmission. Individual differences in vertical transmission, for instance, could be due to when females have been infected by horizontal transmission (Suchman *et al.*, 2006), or whether they have been infected vertically or horizontally. Due to the lack of CpDV free lines, it was not possible to quantify the horizontal transmission or venereal transmission by experimental infections. Nevertheless, the infection rate changed from 17% at the L1 stage to 100% in tested mothers suggesting a high level of horizontal transmission.

The co-existence of endosymbiotic bacteria *Wolbachia* and CpDV within the host ovary cells suggested that they might be interacting. Indeed, symbiotic bacteria and viruses have been shown to interact in several ways. Firstly, symbiotic bacteria can “protect” their hosts against pathogens by either modulating the host immunity or tolerance, or by competing for host resources. A famous example of such interaction is the *Wolbachia* mediated protection of insect hosts against RNA viruses (Hedges *et al.*, 2008; Teixeira *et al.*, 2008). In *Drosophila* this “protection” strongly changed according to *Wolbachia* strains differing in their titers (Martinez *et al.*, 2017). Similar “protection” has been reported in mosquitoes, especially in *Ae. aegypti* transinfected with non-native *Wolbachia* strains from *Drosophila*, against several arboviruses such as Dengue, Chikungunya, West Nile and Yellow fever viruses (Moreira *et al.*, 2009; Mousson *et al.*, 2010; Hussain *et al.*, 2012). On the other hand, native *Wolbachia* in *Ae. albopictus* that exhibit low densities, even in the ovaries, do not influence virus replication (Mousson *et al.*, 2010; Lu *et al.*, 2012). A global pattern both in mosquitoes and *Drosophila* is that *Wolbachia-* mediated antiviral protection seems to depend on *Wolbachia* density (Lu *et al.*, 2012; Osborne *et al.*, 2012; Bian *et al.*, 2013). Our results showed, for the first time, a significant effect of *Wolbachia* strain on the quantity of a DNA virus, the CpDV, in the ovaries suggesting an interaction between these two symbionts sharing the same cells. Indeed by using a backcrossing protocol in mosquitoes naturally infected both with their respective *w*Pip strain and CpDV, we have demonstrated that CpDV density in the ovaries was not controlled by host nuclear genetic background but most likely by maternally transmitted cytoplasmic factors such as *Wolbachia*. However, *Wolbachia* density was really stable between naturally infected *Cx. pipiens* lines for different *Wolbachia* strains tested suggesting variations of symbiont titer was not responsible for variations in CpDV titers. Nevertheless further research including the establishment of CpDV free lines from putative uninfected natural populations followed by experimental infections would be useful to delineate the complex tripartite interactions in this new mosquito-virus-*Wolbachia* natural system.

To further investigate whether CpDV and *Wolbachia* had reciprocal interactions and whether CpDV could still be vertically transmitted in the absence of *Wolbachia*, we treated our lines with antibiotics. This treatment decreased both the amount of CpDV in the ovaries and the vertical transmission alike. However, following the end of the treatment CpDV amount in the ovaries increased in the *Wolbachia*-free lines. This suggested that this sudden decrease could be caused by either the direct influence of antibiotics on the host metabolism, or by the rest of the microbiota that putatively recovered in the generations following the end of the antibiotic treatment. Nevertheless, even though CpDV was able to be transmitted vertically in the absence of *Wolbachia*, the vertical transmission was lower in *Wolbachia*-free lines, suggesting their effect on CpDV transmission. Several endosymbiotic bacteria have been reported to modulate virus transmission. For instance Tomato yellow leaf curl virus’ (a single stranded DNA plant virus) transmission is facilitated by GroEL protein secreted by its vector’s endosymbiotic bacteria *Hamiltonella* (Gottlieb *et al.*, 2010). More recently, Jia et al. (2017) showed that Rice Dwarf virus (a plant double stranded RNA virus) binds to the envelopes of endosymbiotic bacteria *Sulcia* improving its vertical transmission in leafhoppers. Similarly, vertical transmission of *Wolbachia* could be hitchhiked by CpDV in *Cx. pipiens*. CpDV could potentially achieve this through the interactions of its capsids with bacterial surface glycans. Densovirus virions, as of many non-enveloped viruses, are constituted by a single major capsid protein with supersecondary structures which can act as interaction motifs (e.g. Single Jelly Roll) (Neu *et al.*, 2011; Huang *et al.*, 2014). These motifs can attach to bacteria’s surface glycans which can serve as initial attachment factors for many viruses (Neu *et al.*, 2011; Huang *et al.*, 2014). Thus CpDV capsids could potentially interact with a wide array of glycans, including *Wolbachia*’s surface glycans and such interactions could influence the infection outcome and lead to their co-vertical transmission (Kuss et al., 2011; Berger and Mainou, 2018).

Overall we observed a low amount of CpDV in our samples highlighting a covert infection in laboratory *Cx. pipiens* mosquitoes. Viruses exhibiting covert infections can differ in their replication strategy and virulence (Kane and Golovkina, 2010; Williams *et al.*, 2017). They can either cause persistent infections with low levels of virus replication, producing virions but only causing sub-lethal disease or causing latent infections, without any production of virions and limited number of viral gene expression (Kane and Golovkina, 2010; Williams *et al.*, 2017). In the case of CpDV, we were able to observe CpDV virions but not a particularly high mortality in the mosquito lines even in the heavily infected Harash line. This type of covert infection combined with the mixed mode transmission can allow the persistence of CpDV in laboratory as well as natural populations of *Cx. pipiens* mosquitoes. In *Cx. pipiens* mosquitoes even low rate vertical transmission can have a large effect on the epidemiology of CpDV by allowing the transmission of the virus even after acute reduction of host density, e.g. during overwintering diapause (Ebert, 2013; Williams *et al.*, 2017). While still not well studied this type of covert persistent infections by insect specific viruses can largely affect the ecology of the insect populations and vector competence.

## Experimental procedures

### Mosquito Lines

All the mosquito lines used in this study (Supplementary Table 1) have been maintained in insectary conditions (at 25 ± 2 °C and 75 ± 2% relative humidity and a 12:12 h photoperiod). During larval stage, they have been fed with a mixture of shrimp powder and rabbit pellets. Adults have been kept in 65 dm^3^ cages and have been fed with a honey solution. Females were fed weekly with turkey blood using a Hemotek membrane feeding system (Discovery Workshops, United Kingdom).

#### Backcrosses

To test the effect of the *Wolbachia* genetic differences on CpDV amount and their vertical transmission, we introduced the cytoplasm of Tunis and Harash lines, along with their respective *w*Pip strains, into the Slab nuclear background as described by Duron et al (Duron *et al.*, 2012). With this backcrossing protocol we have established BcTunis and BcHarash lines with the same genetic background (over 97% of genome replacement of the original lines by Slab nuclear genome is estimated) and different *Wolbachia* types (*w*PipI and *w*PipIV respectively, Table 1). *w*Pip types were verified with a PCR/RFLP test on pool of larvae using pk1 primers as previously described (Altinli *et al.*, 2018).

#### Wolbachia free lines

BcTunis, Slab and BcHarash lines, all stably infected with wPip (Wo+), were treated with tetracycline (50 mL/L for larval treatment using a 0.4g/L solution) for 3 generations to create *Wolbachia* free lines (Wo-: BCTunisTC, SlabTC, BCHarashTC, Table1). Experiments were conducted using the females from: 1) the first generation right after we started the antibiotic treatment, 2) the third generation during the antibiotic treatment to be able to follow the effect of the treatment and 3) from at least three generations after the end of the antibiotic treatment, to exclude the direct effect of TC treatment on host metabolism, and consequently on CpDV quantity and transmission (Wo-).

### Immunolabelling

Immunolabeling were performed on ovaries from 6 days old females and dechorionated eggs. The egg rafts were collected in 1-4 hours following egg laying. The egg-rafts were bleached (active ingredient, 9.6% of sodium hypochlorite) to separate the individual eggs and rinsed with distilled water. They were then fixed with 3.2% para-formaldehyde diluted in PBS-T (PBS 1X with Tween 0,02%) for 2 hours, and rinsed with PBS 1X. The chorion of each individual egg was removed manually with a needle under an optical microscope (Leica MZ 8).

All the samples were fixed for 1 h in 4% paraformaldehyde (PFA) and washed 3 times with PBS, then treated with 50 mM of NH4Cl for 30 min to minimize autofluorescence. Ovaries and eggs were permeabilized with PBS containing 0.5% Triton and blocking was performed with 1% bovine serum albumin (BSA) in PBS for 1 h. Immunolabelling of CpDV were performed using a mouse primary JcDV anti-capsid antibody that co-reacts with CpDV capsids (Jousset et al., 2000), diluted at 1:300 in PBS with 0.2% Tween and incubated overnight à 4°C. Secondary antibodies (Alexa Fluor^®^ 488 and 568; 1:500; Invitrogen) were used at 1:500 dilutions and incubated for 1 h (ovaries) or 6 h (eggs). Samples were incubated 30 min with Phalloidin-FITC (1:300; Sigma) to label actin cytoskeleton, and 5 min with DAPI (1 μg/ml; Invitrogen) to label nuclei and *Wolbachia*. Finally, labeled tissues and eggs were mounted in Dako (Sigma) on coverslips for observation with a Leica SPE confocal microscope equipped with suitable lasers and bandpass for the dyes (excitation 405 and emission 420-480 for DAPI; excitation 488 and emission 505-550 for Alexa Fluor^®^ 488; emission 532 and emission 580-650 for TexasRed), using a 20X/0.55 HCX APO L W and a 40X/1.15 ACS APO oil objectives. We used the LAS AF (Leica) software to operate the microscope and all images were processed with ImageJ software.

### Electron microscopy

To observe CpDV particles, a pool of ovaries from 6 days old females were dissected, crushed in PBS and filtered with a 0.22 μm pore size filter. The filtered homogenate was subjected to a negative staining using 2% phosphotungstic acid (PTA, Electron Microscopy Sciences) for observation using transmission electron microscopy (TEM, see above).

Ultrathin sections were prepared from ovaries of 6 days old females and dechorionated eggs. Dissected ovaries and eggs were first fixed in 2.5% glutaraldehyde in PBS for 2 h at 4°C, and postfixed in 2% osmium tetroxide in the same buffer for 1 h at room temperature. Samples were then thoroughly rinced in distilled water, stained *en bloc* in 2% aqueous uranyl acetate solution for 2 h at room temperature, dehydrated in increasing-concentration ethanol baths, embedded overnight in Embed 812 resin (Electron Microscopy Sciences), and further polymerized for 2 days at 60°C. Then, ultrathin sections of approximately 100 nm, performed using an LKB Ultrotome III on copper grids, were counterstained with uranyless (Delta Instruments) and lead citrate and examined using a JEOL 1200 EX II transmission electron microscope operating at 100 kV and equipped with an Olympus camera (Quemesa model, 4008 × 2664 pixels) (MEA Platform, University of Montpellier, France).

### Vertical transmission assays

Adult mosquitoes were collected from population cages with a mouth aspirator after 5 days following blood meal. CO2 was used to immobilize the mosquitoes. Ten females from each line (to test the effect of distinct *Wolbachia* strains: Slab, Harash, Tunis, BcTunis, BcHarash; effect of curing *Wolbachia* with tetracycline: SlabTC, BcTunisTC, BcHarashTC, Table 1) were isolated in individual autoclaved glass containers covered with a net. Once females were awake, one third of the glass containers were filled with bottled water that has been tested negative for CpDV prior to use and females were left to lay eggs. Egg-rafts were then transferred to a new sterile glass container and water to hatch, to prevent any possible contamination from females to the water. Five egg-rafts were randomly chosen per line whenever possible and 10 L1 stage larvae from each females have been collected right after they hatched. This was repeated twice adding up to about 10 females and 100 L1 stage larvae per line with a total of 500 L1 tested.

Each L1 stage larva was individually collected to a 1.5 μl Eppendorf tube and crushed in 10 μl of STE buffer (100 mM NaCl, 10 mM Tris-HCl, 1 mM EDTA, pH 8.0) to obtain a crude DNA extraction. Two PCR reactions were conducted using these crude extractions, 1) to check the presence of CpDV using CpDVquantiF and CpDVquantiR primers and 2) to check the amplifiability of DNA using ace-2 gene primers, acequantidir and acequantirev (see above for sequences). The larvae crude extraction that did not allow any amplification in the latter were excluded from further analysis. PCR amplification conditions were as following: i) for CpDVquantiF and CpDVquantiR primers initial denaturation for 5 min at 94 °C, followed by 33 cycles of denaturation, annealing and elongation respectively at 94 °C for 30 s, 56 °C for 30s, and 72 °C for 90 s, and a final elongation at 72 °C for 5 min ii) for ace2quanti primers initial denaturation for 5 min at 94 °C, followed by 35 cycles of denaturation, annealing and elongation respectively at 94 °C for 30 s, 52 °C for 30s, and 72 °C for 90 s, and a final elongation at 72 °C for 5 min.

The proportion of CpDV-positive over CpDV-negative L1 larvae per female is calculated as vertical transmission rate.

### Quantification of CpDV and *Wolbachia*

Ovaries were dissected from 6 days old female mosquitoes. Three ovaries were pooled together and 5 replicates were done per line. Qiagen extraction kit was used to extract DNA of ovary pools according to the manufacturer’s instructions. DNA was eluted in 55 μl PCR gradient water. Negative controls were made to confirm that there was no contamination from the extraction kit and were always negative. Additionally to check the effect of *Wolbachia* and CpDV amount in the females on vertical transmission rate, DNA from the females that laid eggs have been extracted using CTAB technique (Rogers and Bendich, 1989).

Real-time quantitative PCR (qPCR) was used to estimate the amount of *Wolbachia* and CpDV in each sample. qPCR were performed using SYBR Green I Master (Roche) and analysed on a LightCycler 480 Instrument (Roche) using the LightCycler 480 software (version 1.5.1.62). On each DNA sample three PCR reactions were performed in triplicates using i) NS2 protein coding region specific primers to quantify CpDV (CpDVquantiF: 5’-CATTGGAGGAGAAAGTTGGAA-3’, CpDVquantiR: 5’-TCCCCATTATCTTGCTGTCG-3’), ii) wsp specific primers to quantify *Wolbachia* (wolpipdir 5’-AGAATTGACGGCATTGAATA-3’ and wolpiprev 5’-CGTCGTTTTTGTTTAGTTGTG-3’ (Berticat *et al.*, 2002)) and iii) *Culex* Ace-2 locus specific primers to quantify host genome copy as a reference (Acequantidir: 5’-GCAGCACCAGTCCAAGG-3’, Acequantirev: 5’-CTTCACGGCCGTTCAAGTAG-3’) (Weill *et al.*, 2000). Standard curves were created using dilutions of a pCR^®^4-TOPO-Vector (Invitrogen) containing a single copy of each *ace-2, wsp* and *ns2* gene fragments. Plates were designed to have all the compared groups of a biological replicate and prepared using Echo 525 Liquid Handler, Labcyte Inc. The amount of *Wolbachia* and CpDV were normalized with the amount of Ace-2 copies in order to obtain a number of virus or *Wolbachia* per cell.

### Data Analysis

All data analyses have been performed in R version 3.3.1 (R Core Team, 2016).

To analyze the effects of *Wolbachia* amount and diversity, on the amount of CpDV in the ovaries, one highly infected outlier from Harash line has been removed from the analysis and normality of the continuous response variable (log10 transformed amount of CpDV genome/ amount of host genome) has been checked with Shapiro–Wilk normality test. A linear mixed effect model is fitted with genetic background (Slab, Harash, Tunis), *Wolbachia* type (wPip 1, wPip 3 and wPip 4) and *Wolbachia* amount as fixed effects and replicates as random effects using lme4 package (Bates, 2010). Homoscedasticity and normality of the residuals were checked using Breusch-Pagan test (ncvTest, in car package (Fox and Weisberg, 2011)) and Shapiro-Wilk test respectively. Likelihood ratio tests of the full model against the model without a given effect were used to obtain deviance and p-values.

To analyze the effects of *Wolbachia* diversity on vertical transmission of CpDV, a binomial generalized mixed effect model is fitted with genetic background (Slab, Harash, Tunis), *Wolbachia* type (wPip 1, wPip 3 and wPip 4), *Wolbachia* amount and CpDV amount in the full body of the female as fixed effects and replicates as random effects.

CpDV in females were checked for the first and third generation of tetracycline treatment, they were grouped under during tetracycline treatment group given that these two groups didn’t show any significant difference in terms of CpDV amount in the ovaries and represented the same treatment status. A Dunnett’s test (glhrt function in multcomp package) (Hothorn *et al.*, 2008) have been performed for pairwise comparison of treatment stages to the non-treated control group.

In this model genetic background (Slab, Harash, Tunis) and treatment status (no treatment, during TC treatment and after TC treatment) were fixed effects and different replicates are added as random effects. This last dataset also have been analysed using pairwise Wilcoxon test with Bonferroni correction to compare the different treatment stages to the non-treated control group.

## Acknowledgments

We thank Sandra Unal and Patrick Makoundou for technical support. We thank Margareth Capurro for providing us with the “Brasil” *Culex* line. We also thank Dr Philippe Clair for his help in the Real Time Quantitative PCR experiments which were performed through the technical facility of the “qPCR Haut Debit (qPHD) Montpellier genomiX” platform. We acknowledge the imaging facility MRI, member of the national infrastructure France-BioImaging infrastructure supported by the French National Research Agency (ANR-10-INBS-04, «Investments for the future»). Mine Altinli received support from Ministère des affaires étrangères (Campus France) via the French Embassy in Turkey and Project “ISEM Sud”, 2016 for her PhD.

---

**Supplementary Figure 1.**
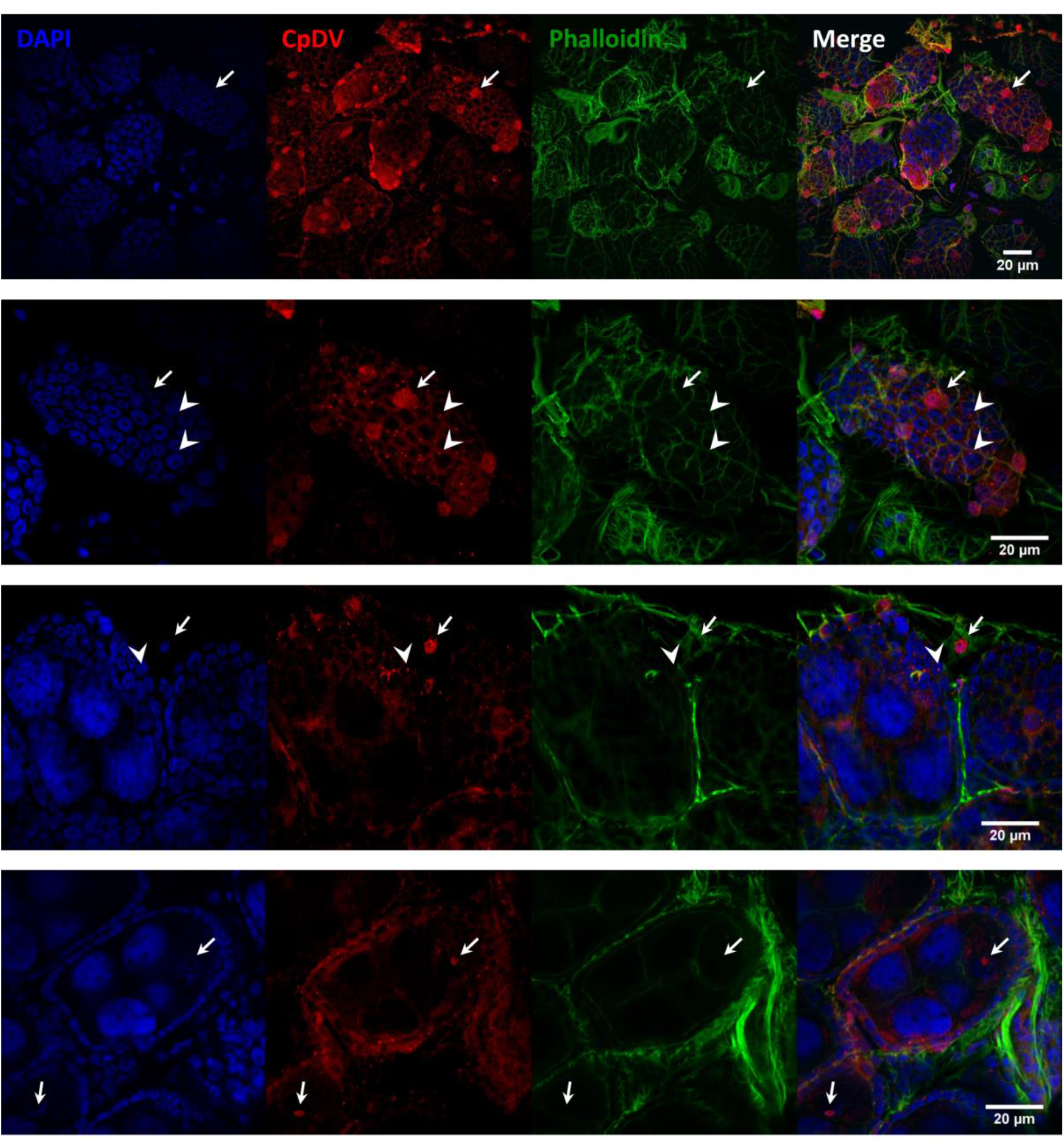

**Supplementary Figure 2.**
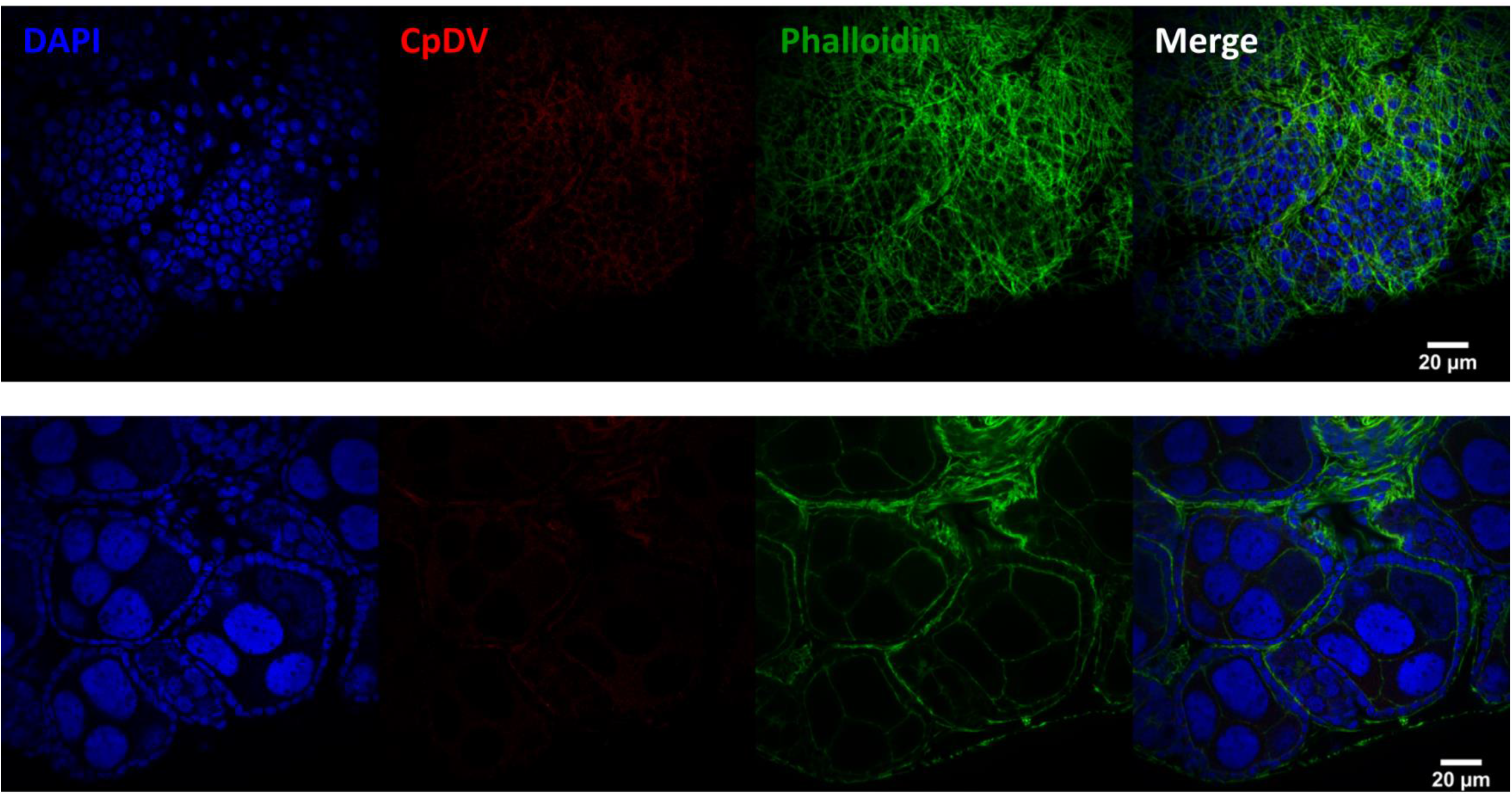

